# Baseline cell proliferation rates and response to UV differ in Lymphoblastoid Cell Lines derived from healthy individuals of extreme constitution types

**DOI:** 10.1101/2021.01.26.428133

**Authors:** Sumita Chakraborty, Sunanda Singhmar, Dayanidhi Singh, Mahua Maulik, Rutuja Patil, Satyam Kumar Agrawal, Anushree Mishra, Madeeha Ghazi, Archana Vats, Vivek T Natarajan, Sanjay Juvekar, Bhavana Prasher, Mitali Mukerji

**Author notes:** Equal contributions (Sumita Chakraborty, Sunanda Singhmar).

## Abstract

Differences in human phenotypes and susceptibility to complex diseases are an outcome of genetic and environmental interactions. This is evident in diseases that progress through a common set of intermediate patho-endophenotypes. Precision medicine aims to delineate the molecular players for individualized and early interventions. Functional studies in Lymphoblastoid Cell Line (LCL) model of phenotypically well characterized healthy individuals can help deconvolute and validate these molecular mechanisms. We developed LCLs from eight healthy individuals belonging to three extreme constitution types, deep phenotyped on the basis of Ayurveda. LCLs were characterized by karyotyping and immunophenotyping. Growth characteristics and response to UV was studied in these LCLs. We observed significant differences in cell proliferation rates between the contrasting groups such that one type (*Kapha*) proliferates significantly slower than the other two (*Vata, Pitta*). In response to UV, one fast growing group (*Vata*) shows higher cell death but recovers its numbers due to inherent higher rates of proliferation. The baseline differences in cell proliferation are key to understanding the survival of cells in UV stress. Variability in baseline cellular phenotypes not only explains the cellular basis of different constitutions types but can also help set priors during designing an individualized therapy with DNA damaging agents. This is the first study of its kind that shows variability of intermediate patho-phenotypes amongst healthy individuals that have implications in precision medicine.

## Introduction

A complex interplay of genetics and environmental conditions lead to inter-individual differences in phenotypic outcomes and disease predispositions ^1,2^. Precision medicine aims to identify genetic variations that help predict these differences ^3^. Strategies are being attempted to uncover variations that govern differential response to the environment in healthy and diseased individuals ^4^,^5^. Identifying variations that govern differential response to the environment in healthy individuals prior to disease onset is still a challenge ^6–8^. This would be important to identify predictive markers for actionable, early and individualized interventions.

Lymphoblastoid Cell Line (LCL) models of genetically characterized individuals such as the 1000 genomes project are being used to identify quantitative trait loci (eQTL) ^9^. In addition, cell lines of diseased individuals are also being used to uncover QTLs linked to differential drug response ^10,11^. However, connecting these cellular differences to phenotypic outcomes require newer and innovative approaches, such as functional phenomics that overlays information from multidimensional phenotypes for understanding biological differences ^12^.

In Ayurgenomics, multi-omics studies are carried out on healthy individuals that are classified on the basis of multi-system phenotypes, called “*Prakrit*i” ^13^. Individuals of a population can be stratified into broadly seven types [10]. *Prakriti* of an individual is invariant and governs health and disease trajectories including responsiveness to the environment. Out of the seven, *Vata, Pitta* and *Kapha* are the most contrasting types in phenotypes and susceptibilities ^14,15^. Unsupervised clustering approaches have revealed distinct phenomic architecture of these contrasting constitutions. Supervised machine learning and advanced statistical models could accurately predict the *Prakriti* types ^13^. These constitution types exhibit differences at the genome wide expression, genetic, epigenetic and microbiome levels ^14,16–19^. Further, *Prakriti* types significantly differ with respect to variations in genes associated with response to double stranded RNA, DNA-damaging agents and hypoxia ^16,20,21^.

These multi-omic differences prompted us to develop cellular models of extreme *Prakriti* types to elucidate differences in cellular phenotypes, which might provide further scope for functional studies. We hypothesized that different *Prakriti* cell lines could respond differently to the same trigger through the same or distinct molecular networks. In this study, we report the development of LCLs derived from healthy humans of extreme *Prakrit*i types, identified from a cohort of a homogenous genetic background ^16^. During LCL characterization, we made a surprising observation-There were striking differences in growth rates amongst the eight LCLs. Intrigued by this observation, we proceeded to establish the robustness of these differences in cell proliferation and their consequence in UV stress. This is the first study that reports a baseline difference in cell proliferation amongst healthy individuals that contributes to variability in response to UV. These observations would be relevant in scenarios where altered rates of cell proliferation are cause or consequence in disease outcomes and help set priors for precision interventions.

## Materials and methods

### Sample collection and ethical clearance

The LCL lines were prepared from healthy individuals of extreme *Prakriti* types identified from the Ayurgenomics cohort from Vadu ^13^. Ten male participants were selected based on detailed clinical phenotyping by Ayurveda clinicians. The study was approved by the KEMHRC Ethics committee. With written informed consent of the participants, 10 ml blood was drawn in heparin vacutainers, stored at room temperature and processed within 24 hours. Blood was tested for Hepatitis B and HIV prior to creation of cell lines. It was ensured that the participants were not under any medication at the time of sampling.

### Peripheral Blood Mononuclear Cells (PBMC) Isolation

Blood was diluted with equal volume of Phosphate Buffer Saline (PBS), mixed by inversion and layered on Ficoll paque density gradient. PBMCs were collected from the buffy layer after centrifugation at 400g for 40 minutes. These were cultured in RPMI-1640 (AT060, Himedia) supplemented with 15% fetal bovine serum (FBS; Gibco, 10270-106). The resultant PBMCs were then used for the generation of LCLs.

### Lymphoblastoid cell line generation

Fresh PBMCs were infected with Epstein - Barr Virus (EBV) and incubated in 15% RPMI at 37°C with 5% CO_2_ in a cell culture incubator for three hours. 1 μg/ml cyclosporine A in 20% RPMI medium was added in 1:1 ratio to the cellular media containing viral particles. Cells were incubated for six days without media supplementation. Subsequently, cells were cultured for three to four weeks with regular media changes. The LCLs were maintained in RPMI-1640 supplemented with 15% FBS, 2 g/L sodium bicarbonate (S5761, SIGMA) and 1X Antibiotic-Antimycotic (15240-062, Gibco). Cells were maintained at 37°C with 5% CO_2_, in a cell culture incubator. LCLs were successfully derived from eight individuals and used in subsequent experiments.

Morphology of the cells with respect to aggregates/clumps and rosette formation was recorded. Cellular clumps were disrupted by pipetting to ensure a multiclonal population. This population was considered to be the first passage of LCLs. Characterization and subsequent experiments were performed from the fifth passage.

### Characterization of LCLs

**(a)Karyotyping**: Karyotyping of LCLs was performed by standard methods.^22^ Briefly, 2 x 10^6^ cells were treated with 10 μl/ml of Colcemid for 30 minutes in normal culture conditions. They were subjected to a hypotonic solution of 0.075M KCl for 30 minutes in a 37°C water bath and fixed in fresh Carnoy’s fixative (3:1, methanol: acetic acid). Slides were prepared by pipetting 2 drops of cell suspension on slides held at 45 degrees. Slides were heat fixed on a 45°C hot plate for 2 minutes. Slides were aged for a day followed by trypsinization and staining with Giemsac stain for visualizing the chromosome banding pattern [1]. **(b)Immunophenotyping**: In order to ensure that the LCLs originated primarily from B cells, immunophenotyping was carried out by flow cytometry analysis of B cell specific surface markers. 0.1×10^6^ cells were harvested and washed with PBS for surface staining with lymphocyte specific CD19 (B cell), CD3 (T cell) and CD56 (NK cell) markers. PBMCs with heterogeneous cells were used as positive controls. Non-specific binding of the antibody was blocked using FC receptor blocker for 30 minutes on ice followed by incubation with the antibodies for 40 minutes on ice. Cells were washed with cold PBS twice and immediately analyzed by flow cytometry on BD LSR II. A minimum of 1×10^4^ cells were recorded per sample for all flow cytometry analysis.

### LCL growth curve assay

Growth assay of LCLs was performed using a non-lytic, luminescence based method using the RealTime-Glo™ MT cell viability assay (G9711, Promega). The Realtime-Glo™ enzyme and substrate were added to the cell culture medium during seeding of cells. Cells were seeded at 10000/100μl media per well in a 96-well plate and incubated in normal culture conditions. Luminescence units were recorded at near regular hours for plotting the growth curves. Growth rate and doubling times were calculated from the log phase.

### CFSE proliferation assay

Cell proliferation rates were estimated by tracing cells labelled with Carboxyfluorescein Succinimidyl Ester (CFSE) across generations and compared between populations using CellTrace™ CFSE Cell Proliferation Kit (C34554 Invitrogen) ^23^. Cells were stained with 5 μM CFSE and grown in 12 well plates at the seeding density of 1×10^5^/ml. Readings were taken on day zero and day 3 (72 hours). Cells fixed in 4% Paraformaldehyde were resuspended in FACs buffer (PBS with 1% FBS) and analyzed by FACS on BD LSR II. The Division Index (DI) - the average number of cell divisions that a cell in the original population has undergone was calculated using FlowJo software (BD).

### Cell cycle analysis after synchronization by double thymidine block

Cells were seeded at the density of 1×10^5^/ml in media supplemented with 2mM Thymidine (T9250-5G Sigma) followed by 18 hours of growth. Cells were washed and grown in media without Thymidine for 9 hours, followed by a 15-hour growth period in Thymidine supplemented media. Cells were harvested and fixed in 70% ethanol followed by permeabilized in 0.1% triton X 100 and RNase(40ng/ml) treatment. Cell cycle phases were studied by staining the cells with Propidium Iodide and analyzing by flow cytometry.

### Comparison of cell cycle stages using specific markers

The proportion of cells in different cell cycle phases were compared by Western blotting. We checked CDK2 Tyr15 phosphorylation for G1/S phase and Histone 3 Ser10 phosphorylation for mitosis. Cell lysates were prepared at 24 hours of growth and SDS-PAGE performed on 15% bis-acrylamide gel. Proteins were electrophoresed onto a PVDF membrane and blocked with 5% Bovine Serum Albumin. The primary antibody used was a cocktail of rabbit anti-Cdk2 pTyr15, and rabbit anti-Histone H3 pSer10 (1:1000 dilution, ab136810, abcam). Secondary antibody was horseradish peroxidase conjugated goat anti-rabbit immunoglobulin G (1:10000 dilution, sc-2004; Santa Cruz Biotechnology). Membranes were developed with optiblot ECL detection kit (ab133406; Abcam) and imaged using a Chemi-Doc (ImageQuant LAS 500). Protein expression was measured by densitometry analysis of the bands using ImageJ and normalized to the density of the ß-actin.

### Ki67 assay

To estimate the number of proliferating cells, we carried out a 48-hour time course assay of Ki67. Cells were cultured at 1 x 10^5^ cells/ml in 12-well plates, harvesting one well of each LCL every 12 hours, from 0 to 48 hours. Cells were fixed in 4% Paraformaldehyde, permeabilized with 0.1% Triton x 100 and stained with 1:200 dilution of Ki67 antibody (11-5698-80; Invitrogen) for 40 minutes at RT. Cells were resuspended in the FACS buffer and analyzed on BD LSR II flow cytometer.

### Cell Proliferation and death in response to UV

1×10^5^ CFSE stained cells were seeded per well in 12 well plates. Cells were exposed to 50 mJ/cm^2^ UV B and grown for 48 hours. Live and dead cells were counted using trypan blue staining. Cell proliferation was quantitated by flow cytometry analysis of CFSE stained cells.

### Statistical analysis

Normal distribution was checked using Shapiro-Wilk test. Pairwise comparisons of *Vata* versus *Kapha, Vata* versus *Pitta* and *Pitta* versus *Kapha* were carried out using Student’s t-test.

## Results

### Characterization of *Prakriti* specific LCLs

LCLs showed hallmark morphology with presence of uropods and formation of aggregates in suspension culture, as seen by bright field microscopy (Fig.S1). Karyotype analysis confirmed the presence of normal chromosome numbers, structure and ploidy (Fig.S2). Immunophenotyping affirmed LCLs to be of B cell origin, based on CD19 positive staining (Fig.S3). Cells did not stain for the markers of T cells (CD3) and NK cells (CD56).

### Differences in growth rate between *Prakriti* LCLs

Our initial aim was to study cellular response to UV amongst LCLs for which we needed to work with equal cell densities. However, during initial experiments we observed the *Vata* and *Pitta* cell lines to grow faster in comparison to *Kapha* and requiring more frequent media changes. This prompted us to explore whether there were inherent differences in cell proliferation rates among the LCLs. Growth curve of LCLs revealed two groups of slow and fast growing lines based on the average doubling time (36.6 hours). Interestingly, the three *Kapha* LCLs had doubling times greater than 36.6 hours and fell into the slow growing group (Fig.1A, B; Fig.S4). The three *Vata* and two *Pitta* LCLs had doubling-times less than 36.6 hours and fell into the fast growing group (Fig. 1A, B; Fig.S4). These differences significantly differentiated *Kapha* from *Vata* and *Pitta* (Fig.1A, B). The growth curves were repeated multiple times as biological and technical replicates i.e., a new stock from the same passage was revived for each experiment and experiment was performed in triplicates.

**Figure 1.**
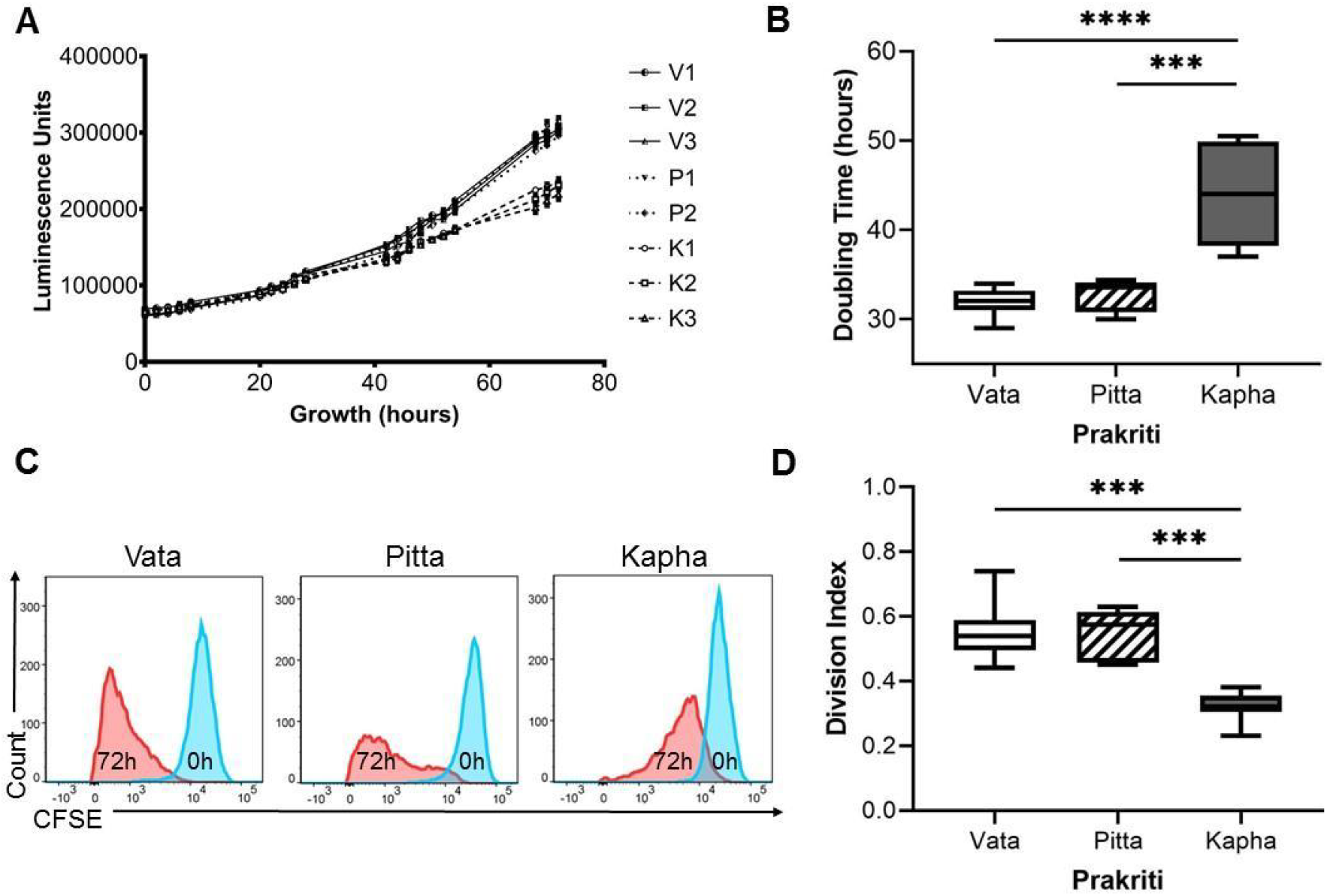
*Prakriti* specific growth differences in LCLs (A) RealTime-Glo™ growth curves of LCLs (B) Doubling times of the *Kapha (K*) to be significantly lower than *Vata* (*V*) and *Pitta* (*P*). Growth curves and box plots from biological and technical replicates [N=3 (*Vata* and *Kapha*) and N=2 (*Pitta*), n=3, Error bars denote ±SEM, unpaired t test **** = P<0.00001, *** = P<0.0001)] (C) CFSE assay shows *Kapha* cell population dividing slower than *Vata* and *Pitta* (representative images of flow cytometry) (D) DI of *Kapha* is significantly lower than *Vata* and *Pitta*. Box plot from analysis of biological and technical replicates [N=3 (*Vata* and *Kapha*) and N=2 (*Pitta*), n=3, Error bars denote ±SEM, unpaired t test *** = P<0.0001)]

### Proliferation differences in cell populations of *Prakriti* LCLs

To investigate whether differences in growth rate is due to the entire population or only a certain fraction, we used the CFSE dye dilution method. We looked at the distribution of the cells across generations at 72 hours of growth. *Vata* and *Pitta* LCLs were almost in the third generation of division whereas *Kapha* LCLs were in the second generation (Fig.1C, S5). We did not see any population of cells in the undivided peak, at 72 hours, which shows that all the cells in the population have undergone at least one round of division. The DI of *Kapha* LCLs is significantly lower than *Vata* and *Pitta* LCLs (Fig.1D).

### Cell cycle kinetics differ between *Prakriti* LCLs

To compare cell cycle kinetics, we planned to synchronize the eight LCLs and follow the progression of cell populations through G1, S and G2 phase by Propidium Iodide staining of the DNA. We used a double thymidine block which arrests cells at the boundary of G1/S. We failed to get the same degree of synchronization in all eight cell lines (S6). We also tried other methods, eg. Serum starvation for 24 hours, nocodazole treatment etc. but failed to get all the LCLs synchronized to the same phase. In an alternate strategy, we used cell cycle marker proteins CDK2 (G1/S) and Histone 3 (mitosis) for inferring differences in kinetics. The G1/S phase marker CDK2 phosphorylation was significantly different between the three groups with levels in *Vata<Pitta<Kapha* at 24 hours of growth (Fig.2A, B). On the contrary, the mitosis marker Histone 3 phosphorylation on the other hand was significantly different between the three LCL groups with levels in *Kapha<Vata<Pitta* (Fig.2A,C).Thus, our results indicates that at 24 hours of growth, mitosis is lower in *Kapha* cell lines compared to others. However, it is interesting to note that even though *Vata* and *Pitta* show higher cell proliferation rates, the molecular mechanisms could be different.

**Figure 2.**
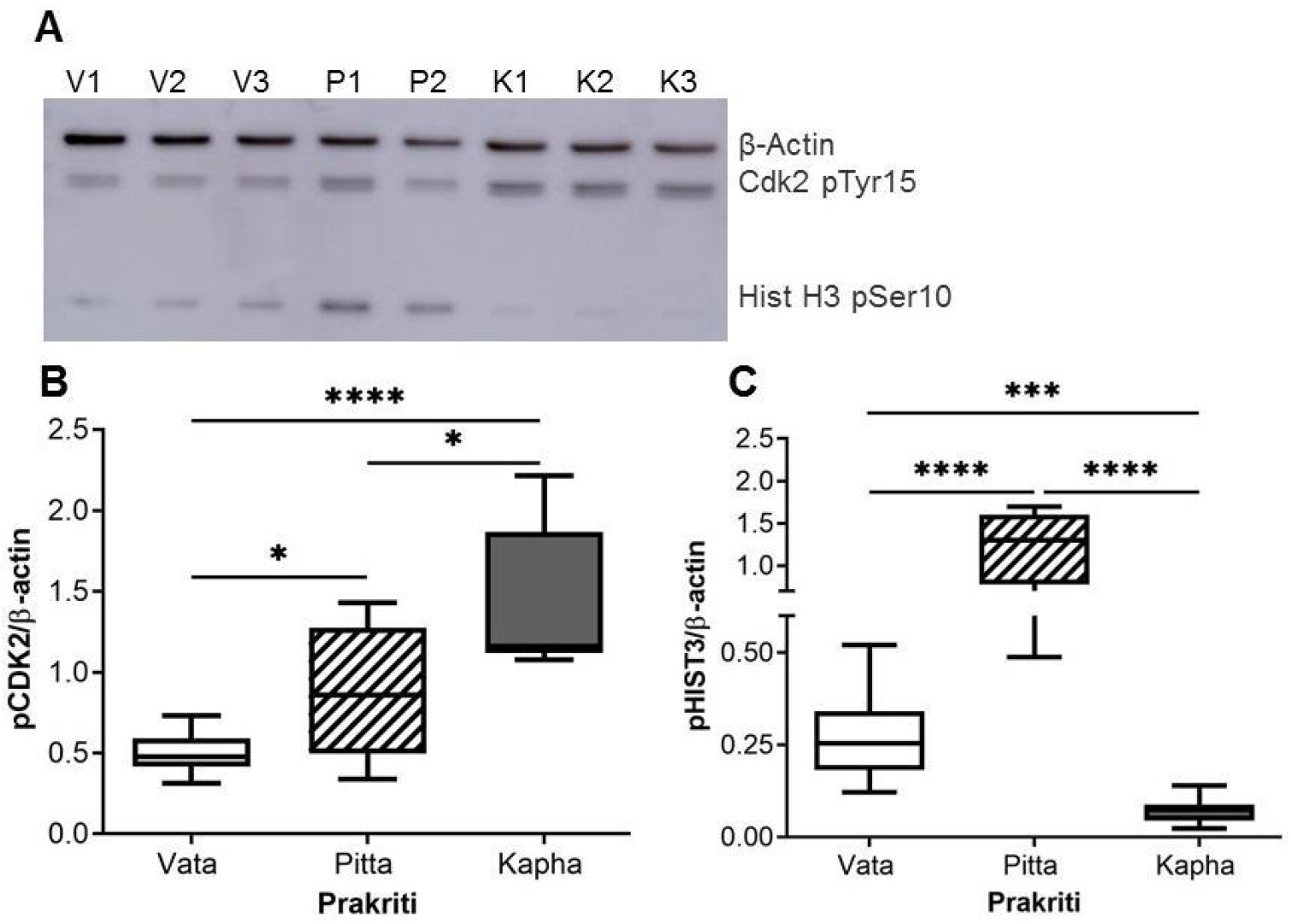
Cell cycle marker proteins in *Prakriti* specific LCLs (A) Western blot (representative) for CDK2 pTyr15 (G2/S marker), Histone H3 pSer10 (Mitosis marker) and β-actin (loading control) at 24 hours of growth (B) Western Blot quantitation showing a significant difference in CDK2 pTyr15 (G2/S marker) between *Prakriti* (C) Histone H3 pSer10 (Mitosis marker) levels significantly differ between *Prakriti* [N=3 (*Vata* and *Kapha*) and N=2 (*Pitta*), n=3, Error bars denote ±SEM, unpaired t test **** = P<0.00001, *** = P<0.0001, * = P<0.05]

To further explore whether the number of proliferating cells is lower in *Kapha* LCLs as compared to *Vata* and *Pitta*, we traced the cell proliferation marker Ki67 in the cell lines over a 48 hour time course. The levels of Ki67 positive cells were higher in the *Vata* and *Pitta* LCLs along the entire time course (Fig.3B) however, significant differences were observed at the 12 hours (Fig.3A, C).

**Figure 3.**
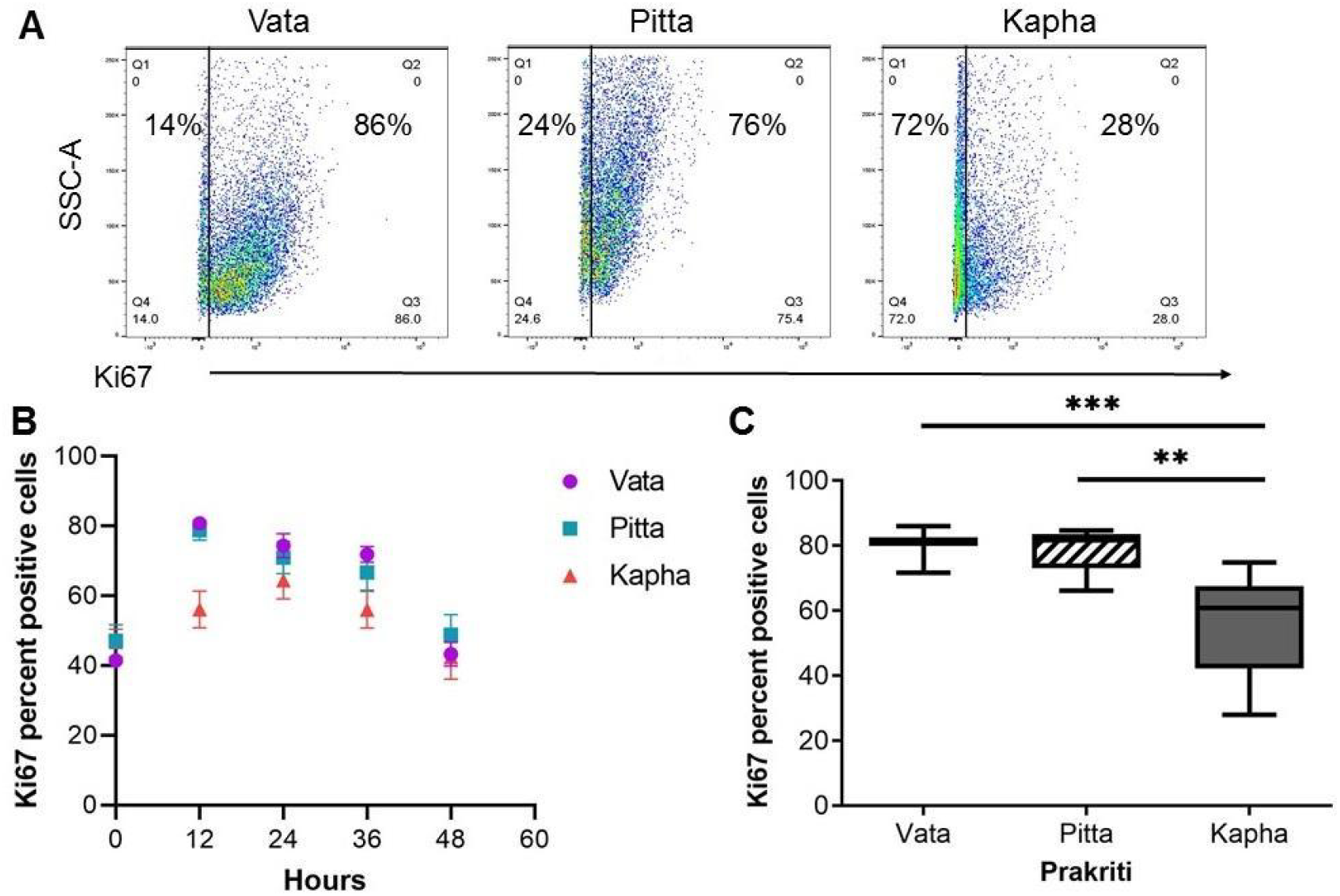
Ki67 proliferation marker differs between *Prakriti* types (A) Percent of proliferating cells in LCL populations at 12hr (Representative flow cytometry plots) (B) Ki67 staining over the course of 48 hours of growth. (C) *Kapha* has significantly lower Ki67 positive cells at 12 hours compared to *Vata* and *Pitta*. [N=3 (*Vata* and *Kapha*) and N=2 (*Pitta*), n=3, Error bars denote ±SEM, unpaired t test *** = P<0.0001, ** = P<0.001)

### Effect of UV exposure on cell proliferation rates in *Prakriti* LCLs

It is known that cell cycle arrest ensues in response to UV or DNA damaging agents. We were curious to know how the LCLs of different *Prakriti* types, with inherent differences in baseline cell proliferation rates respond to UV. Interestingly we found the *Vata* LCLs to be significantly more sensitive to UV than others (Fig.4). They not only show significantly less percentage viability compared to *Kapha* (Fig 4.A), but also a significantly higher death percent compared to both *Pitta* and *Kapha* (Fig 4.B) in response to UV. The *Kapha* LCLs displayed significantly lower growth arrest compared to *Vata* LCLs (Figure 4 E) 48-hours after UV exposure. *Pitta* LCLs did not show any significant differences compared to *Vata* but exhibited higher growth arrest compared to *Kapha* LCLs (Fig 4.E) in UV stress. All the LCLs show similar viable cell numbers after 48 hours of UV exposure (Fig 4.B). But the differences in UV response can only be appreciated by accounting for the baseline viability difference among them (Fig 4.B). All the LCLs show growth arrest in response to UV, but the growth rate of *Vata* still remains higher than the *Kapha* cell lines (Fig.4F). This high growth rate helps the *Vata* LCLs to cope up with the higher cell death caused by UV (Fig.4C, D) and recover their cell numbers (Fig.4B).

**Figure 4.**
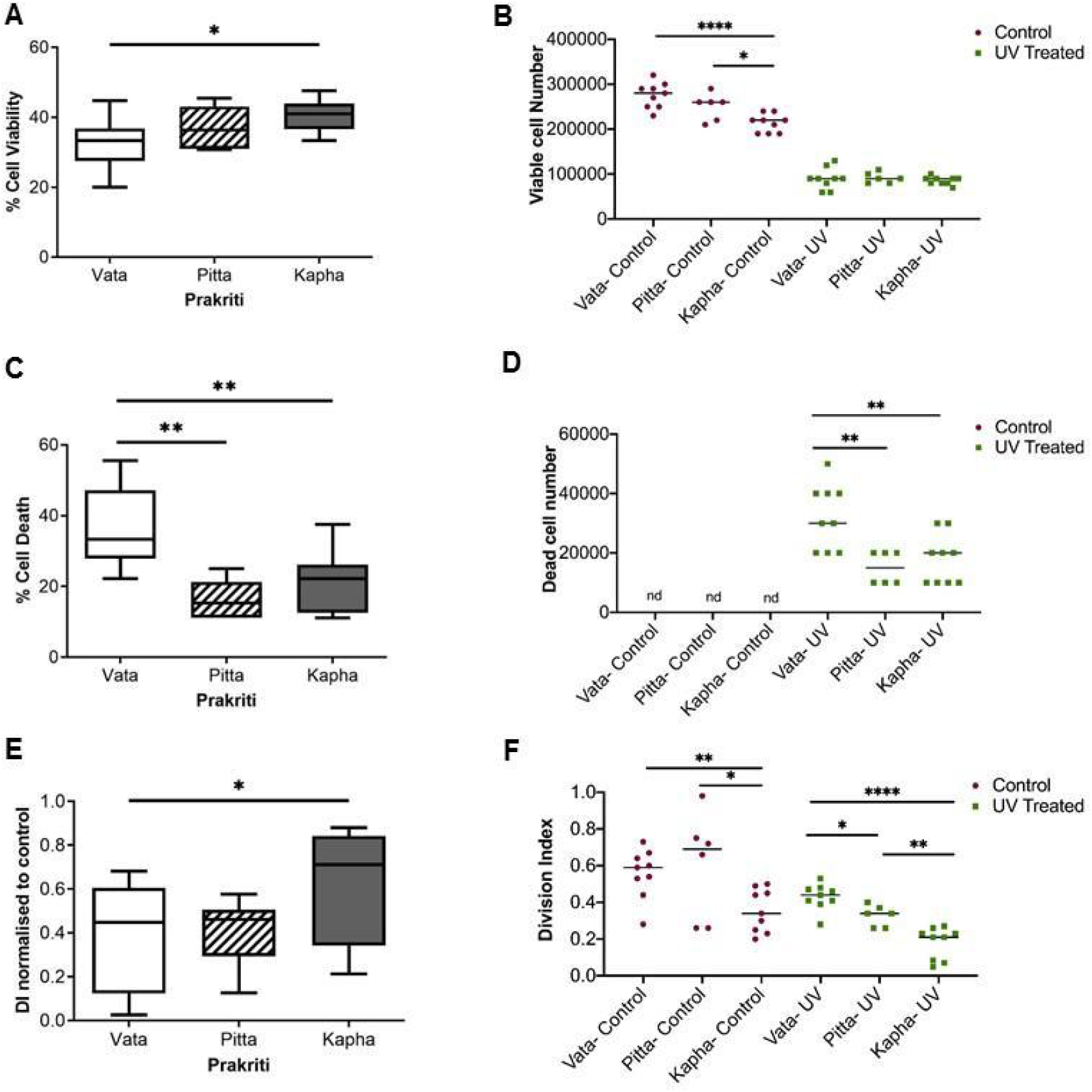
Effect of UV exposure on *Prakriti* specific LCLs. (A) Percent cell viability, (B)Viable cell number, (C) Percent cell death, and (D) Dead cell number of *Prakriti* specific LCL types in response to UV stress. (E) Division index of *Prakriti* LCLs normalized to respective untreated controls (F) Division index of control and UV exposed LCLs. (N=3 (*Vata* and *Kapha*) and N=2 (*Pitta*), n=3, Error bars denote ±SEM, unpaired t test **** = P<0.00001** = P<0.001, * = P<0.05)

## Discussion

We report a novel finding that cell proliferation rates differ significantly amongst LCLs derived from healthy individuals stratified phenotypically. Processes such as cell proliferation are thought to be homeostatic and not expected to vary systematically across individuals, our data suggests otherwise. The LCLs studied here were derived from individuals of extreme and contrasting constitutions who differ with respect to multisystem and molecular attributes ^16–19,24^ We show a core cellular phenotype differentiating between these groups where *Vata* and *Pitta* have higher rates of cell proliferation compared to *Kapha*. However, *Vata* shows significantly more cell death in UV, than *Pitta* and *Kapha*. Despite its higher cell mortality, the baseline proliferation rates in *Vata* seem to provide a growth advantage to the surviving cells post UV treatment.

Cell proliferation is coupled to a variety of cellular and individual phenotypes for instance, DNA repair ^25,26^. Slow proliferation allows repair to be complete before the next cell division whereas fast proliferation allows coupling of DNA repair to replication and enables homology driven replication coupled repair process ^27^. Impaired cell turnover can lead to extensive cell accumulation, e.g. neoplastic and cancerous growths. Inappropriately triggered cell death can lead to excessive cell loss, e.g. neurodegenerative, autoimmune and cardiovascular disorder ^28,29^. Interestingly, neurons re-enter cell cycles before apoptosis and impaired cell cycle regulation is a clear indicator of neurodegenerative disorders ^30^. Therefore, inherent differences in cell proliferation and repair in healthy states could contribute to inter-individual variability in disease and responsiveness. In this context, a recent study reports higher basal cell proliferation rate differences in LCLs derived from patients of Bipolar disorder who are responders to Lithium chloride. The response of control and patient LCLs to LiCl is shown to be dependent on these baseline proliferation differences ^31^. Higher cell proliferation has also been reported in lymphocyte lines of Alzheimer’s disease ^32^. Could this be an early phenotype of predispositions? Assessment of cell proliferation also forms the basis for therapeutic interventions in cancer and its therapies. Without a prior knowledge of baseline proliferation rates, enhanced killing of cancer cells by therapy might suggest a need for lower dosage or shorter treatment. However, priors set from the baseline data may suggest a higher dose or recurrent treatment to prevent relapse or predict differences in responsiveness.

The *Prakriti* types are an outcome of relative proportions of the three groups of physiological entities “*Tridosha*”, that govern the kinetic “*Vata*”, metabolic “*Pitta*” and “*Kapha*” structural components of the body. These proportions in *Prakriti* types, is established at the time of embryonic development, and remains invariant throughout lifetime ^14,24^. Our previous report on the blood transcriptome from another cohort shows differential enrichment of molecular pathways between the *Prakriti* types ^16^. Significant upregulation and enrichment of cell cycle genes between *Vata* and *Kapha* corroborates the cellular differences reported here. We also observed significant genetic differences in a subset of genes which govern cell proliferation, DNA damage response, apoptosis - *AKT3, FAS, RAD51* between the *Prakriti* ^20,33^. Hence, cellular proliferation rate seems to be a core cellular phenotype specific to *Prakriti*. It is also a shared endophenotype for multiple triggers. Deconvolution of this pathway through *Prakriti* phenotypes might provide novel biological leads. For instance, we observe differences amongst *Prakriti* types in phosphorylation of CDK2, a candidate gene implicated in neuronal loss in Parkinson’s disease ^34^.

The observations made here might help provide clues to some of the biological differences and inherent susceptibilities described for *Prakriti*. Noteworthy to mention is the barrier function differences with respect to UV that have been described in many studies ^35,36^. Membrane activating agents have been shown to activate cell division ^37^. In Ayurveda, there are vivid descriptions about better barrier functions in *Kapha*. It might be possible that fast proliferation doesn’t let the cell membrane stabilize as much as the slow proliferators, resulting in poor barrier function. Earlier we have reported such links between hypoxia response and hemostatic outcomes in *Prakriti* types. Individual’s bleeding outcomes were due to inherently elevated hypoxia responsiveness in *Pitta* ^21^. In future studies, it is pertinent to explore whether differences in cell membrane properties could explain the response to UV in *Kapha* and consequent differences in cell proliferation rates.

These observations are preliminary and have to be carried out in larger sample sizes, cohorts and cellular models for generalizability. LCLs have inherent limitations of not being readily amenable to transfection and therefore differentiated cell lineages from the same individual can be considered. Correlation of this core cellular phenotype across different pathological conditions in prospective cohorts is testable. This study also emphasizes the advantage of *Prakriti* based stratification for predictive medicine and early interventional possibilities.

## Conclusion

This first of its kind study highlights the importance of Ayurgenomics approach in the development of cellular models for functional phenomics. These can be effective in integrating the expression readouts from system level phenotypes to cellular outcomes in an individual specific manner. Our observations open up tantalizing possibilities in precision medicine, for instance, taking into account the baselines of the cell proliferations as priors might help resolve unexplainable outcomes and lead to individualized interventions. Non-invasive *Prakriti* based stratification that focuses on individuality is what is envisaged in affordable precision medicine initiatives.

## Acknowledgment

We thank Odity Mukherjee for helping with the LCL facility, and the CSIR FACS Facility. This work was financially supported by the Ministry of AYUSH center of excellence (GAP0183) and CSIR funding by Trisutra (MLP901). Fellowship support was received from CSIR (Sumita), ICMR (Sunanda) and COE (Dayanidhi). We thank the field team of KEM Hospital Research Centre, Pune for assistance in data generation. Contribution of the participants is appreciated.

