## Supplementary Figures for "Baseline cell proliferation rates and response to UV differ in Lymphoblastoid Cell Lines derived from healthy individuals of extreme constitution types"

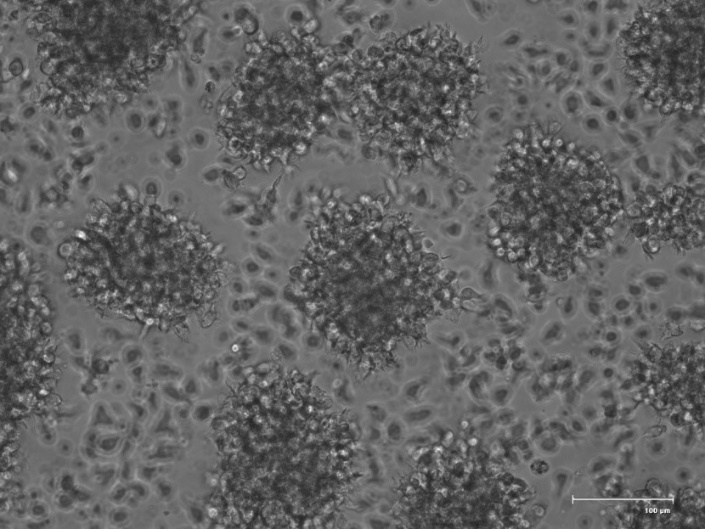

S1. Morphology of LCLs- Cells shows the formation of clumps in suspension culture known as rosette morphology. Representative bright field image, scale bar - 100µm.

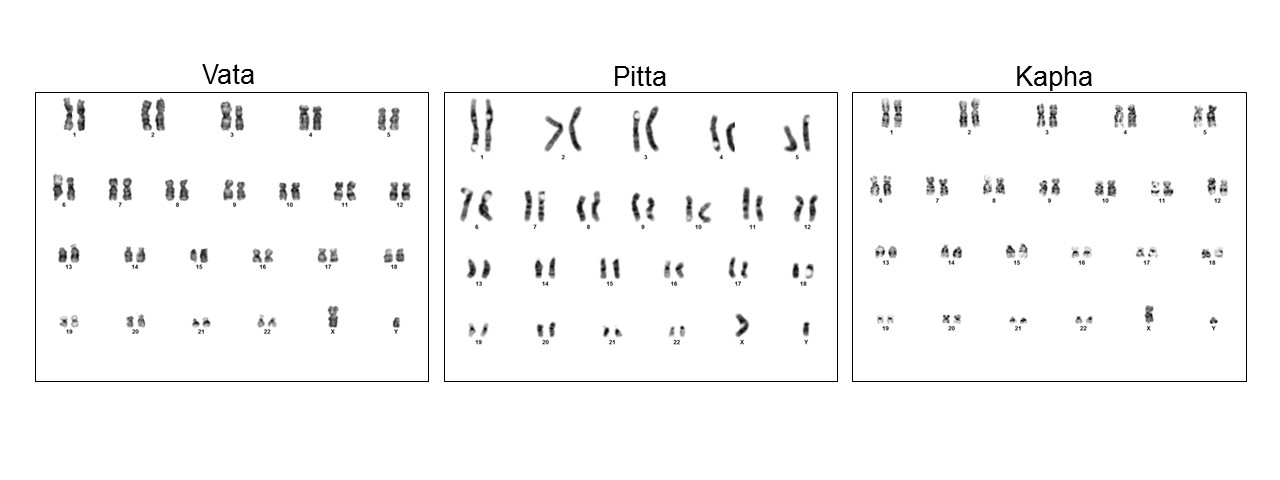
S2. Karyotype analysis of the LCLs- Chromosome of the eight LCLs were analyzed by Geimsa banding and showed diploid, XY karyotype without detectable aberrations. (Representative images of one LCL from each *Prakriti* type).

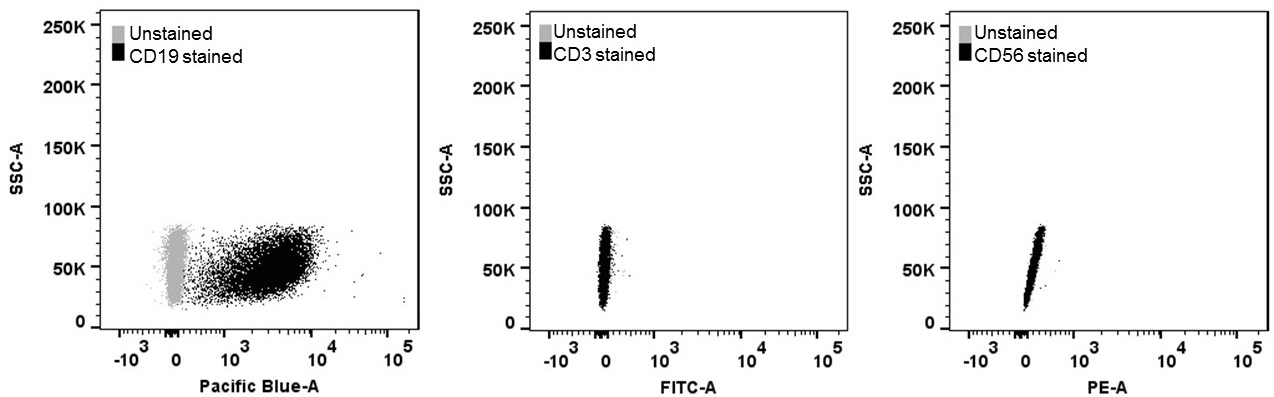

S3. LCLs have B-cell origin- LCLs stained positive for marker of B cells (CD19), and negative for T cells (CD3) and NK cells (CD56) as analyzed by flow cytometry. PBMCs were used as positive control for antibodies. (Representative image).

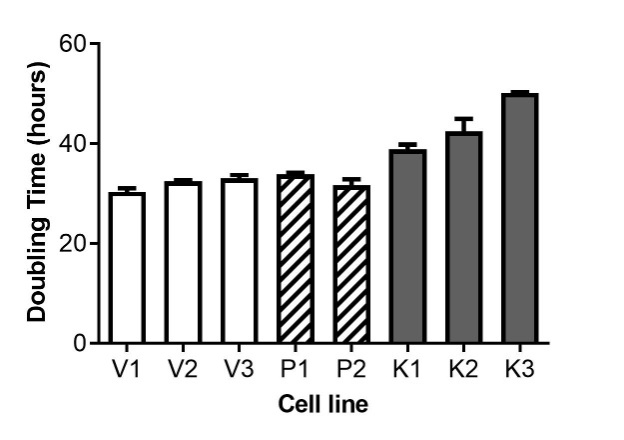

S4. LCL doubling times are in the range of 30 to 50 hours. *Kapha* LCLs have doubling-time greater than the average 36.6 hours (slow growing group). *Vata* and *Pitta* LCLs have doubling time less than 36.6 hours (fast growing group). Error bars denote ±SEM from three biological and three technical replicates.

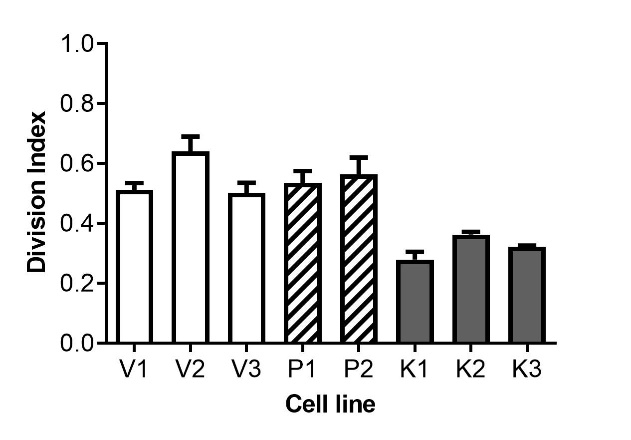

S5. Division index (DI) of LCL cell populations. *Kapha* LCLs have lower DI than *Vata* and *Pitta* LCLs Error bars denote ±SEM from three biological and three technical replicates.

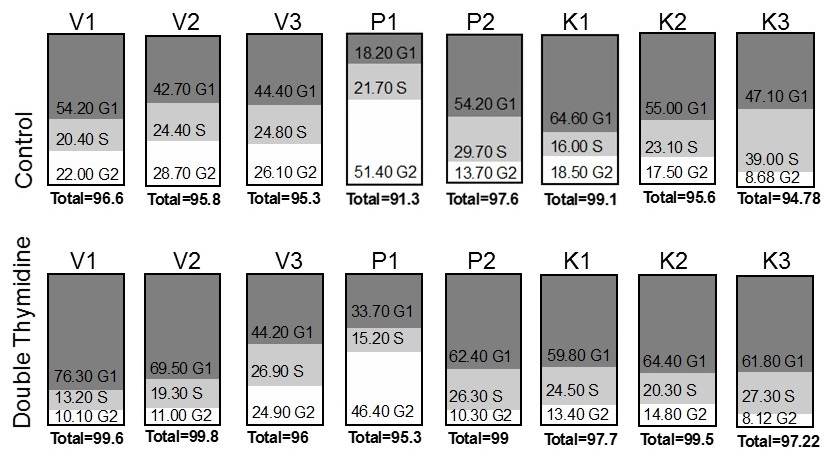

S6. *Prakriti* LCLs synchronized at the boundary of G1/S by double Thymidine block. LCLs show variable synchronization.

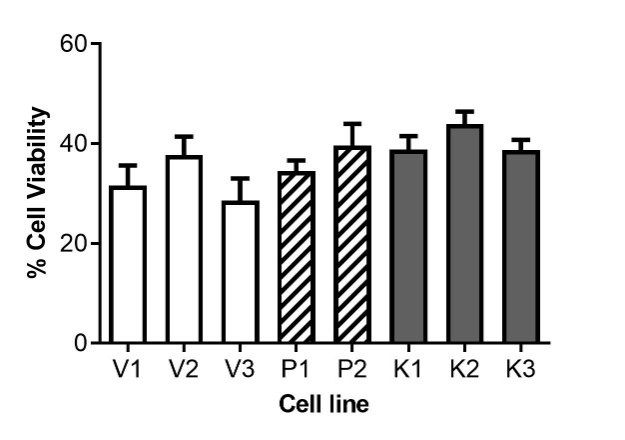

S7. Cell viability after 48 hours of UV exposure. Percent cell viability calculated with respect to untreated controls by cell counting on hemocytometer. (Error bars denote ±SEM from three biological and three technical replicates).

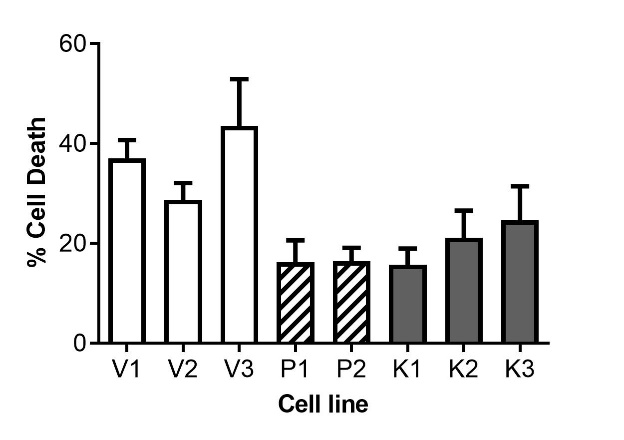

S8. Cell death after 48 hours of UV exposure. Percent cell death was calculated with respect to total number of cells at 48 hours in that sample. Dead cells were enumerated by counting trypan blue positive cells on hemocytometer. (Error bars denote ±SEM from three biological and three technical replicates).

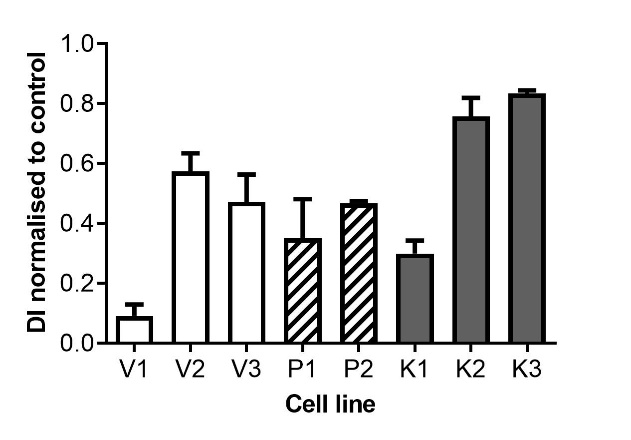

S9. Growth of UV exposed LCLs assessed by Division index (DI) normalized to untreated controls at 48 hours of growth.
